# Mechanisms of frustrated phagocytic spreading of human neutrophils on antibody-coated surfaces

**DOI:** 10.1101/2022.02.18.481104

**Authors:** Emmet Francis, Hugh Xiao, Lay Heng Teng, Volkmar Heinrich

## Abstract

Complex motions of immune cells are an integral part of diapedesis, chemotaxis, phagocytosis, and other vital processes. To better understand how immune cells execute such motions, we present a detailed analysis of phagocytic spreading of human neutrophils on flat surfaces functionalized with different densities of immunoglobulin G (IgG) antibodies. We visualize the cell-substrate contact region at high resolution and without labels using reflection interference contrast microscopy (RICM) and quantify how the area, shape, and position of the contact region evolves over time. We find that the likelihood of the cell commitment to spreading strongly depends on the surface density of IgG, but the rate at which the substrate-contact area of spreading cells increases does not. Validated by a theoretical companion study, our results resolve controversial notions about the mechanisms controlling cell spreading, establishing that active forces generated by the cytoskeleton rather than cell-substrate adhesion primarily drive cellular protrusion. Adhesion, on the other hand, aids phagocytic spreading by regulating the cell commitment to spreading, the maximum cell-substrate contact area, and the directional movement of the contact region.

**Summary:** The detailed analysis of immune-cell spreading on antibody-coated surfaces establishes that active cytoskeletal protrusion rather than passive substrate adhesion drives phagocytic spreading.

## Introduction

Innate immune cells, in particular white blood cells, stand out among animal cells due to their deformability and extraordinary motility. Their mechanical flexibility enables them to carry out vital immune functions, such as chemotaxis toward sites of infection, and phagocytosis of foreign particles. These and other processes critically depend on the cells’ ability to spread onto surfaces, for example, during firm arrest of white blood cells at the endothelium, or during the engulfment of target particles.

Phagocytic spreading is distinct from other forms of cell spreading in that it engages specific cell-surface receptors known to trigger phagocytosis (Cougoule et al., 2004; Flannagan et al., 2012; Lim et al., 2017). In many physiological situations, phagocytosis is initiated by opsonic ligands such as immunoglobulin G (IgG) antibodies and complement fragments. When bound to a pathogen surface, these ligands are recognized by the immune cells’ Fcγ receptors (FcγRs) and complement receptors, respectively (Bournazos et al., 2016; Dustin, 2016). Ligation of these receptors activates receptor-specific intracellular signaling (Flannagan et al., 2012; Futosi et al., 2013; Rosales and Uribe-Querol, 2017; Uribe-Querol and Rosales, 2020) and ultimately leads to progressive spreading of the phagocyte over the target surface. In conventional phagocytosis, cell spreading concludes when the phagocytic cup closes to create a phagosome compartment. However, in some cases known as frustrated phagocytosis, the target is too large to be completely engulfed by a single phagocyte. During these encounters, immune cells strive to maximize the area of contact with the target surface, resulting in thinly spread, lamella-shaped cells that partially cover the target. Such frustrated phagocytic spreading has been observed, for instance, in interactions of human neutrophils with spherules of the fungal pathogen *Coccidioides posadasii* (Lee et al., 2015) as well as with large beads (Herant et al., 2005). The extent of spreading in these cases seems to be limited by the maximum apparent cell-surface area that a phagocyte can generate. For example, the maximum surface area observed during phagocytosis by human neutrophils amounts to about 300% of the surface area of the spherical resting shape of these cells (Hallett and Dewitt, 2007; Herant et al., 2005; Lee et al., 2015; Simon and Schmid-Schonbein, 1988), while for macrophages this plateau value is as high as 600% of the respective resting area (Cannon and Swanson, 1992; Lam et al., 2009).

The targets encountered by immune cells during frustrated phagocytosis are characterized by very low surface curvatures. Although curvature-dependent variations in phagocytic behavior have been reported (Cannon and Swanson, 1992; Champion and Mitragotri, 2006; Pacheco et al., 2013; Paul et al., 2013; Simon and Schmid-Schonbein, 1988), they do not appear to reflect qualitative changes in the cellular response but are likely due to the same response program coping with different “input” sizes and shapes of target particles (Herant et al., 2006; van Zon et al., 2009). Thus, it is reasonable to assume that the same preprogrammed cellular response also controls phagocytic spreading of immune cells on surfaces with zero curvature, i.e., flat substrates. Considering the high resolution of imaging methods that inspect cell-substrate interactions directly at the bottom of an experiment chamber, the study of frustrated phagocytic spreading of immune cells on such planar surfaces is a powerful tool to examine fundamental mechanisms of phagocytosis (Barger et al., 2019; Jaumouille et al., 2019; Kovari et al., 2016; Ostrowski et al., 2019).

Here, we present an in-depth quantitative analysis of human neutrophil spreading on IgG-coated coverslips. Our main objective is to address the lacking consensus between two different hypotheses that have been put forward to explain the mechanistic origin of phagocytic spreading, denoted as “Brownian zipper” and “protrusive zipper” mechanisms (Heinrich, 2015). Both hypotheses agree that fresh contact between the phagocyte membrane and target surface results in a zipper-like adhesive attachment that is essentially irreversible. However, they differ in their assumptions about the primary driving force of cellular protrusion that produces fresh cell-target contact in the first place. The Brownian zipper hypothesis views the phagocyte as a passive object and postulates that strong adhesion alone is responsible for pulling the cell membrane onto the target surface. In other words, this hypothesis treats phagocytic spreading as a wetting phenomenon, akin to a water droplet spreading on a hydrophilic surface. In contrast, the protrusive zipper hypothesis assumes that the phagocyte’s cytoskeleton actively generates protrusive force to push the front of the cell outwards.

We investigate which of these hypotheses captures the main driving force of phagocytic spreading using well-controlled frustrated phagocytosis experiments. Our strategy is based on the following reasoning. If the Brownian zipper hypothesis of adhesion-driven spreading holds true, then both the rate as well as the extent of spreading should strongly depend on the density of adhesive binding sites on the substrate. In contrast, because the protrusive zipper hypothesis assumes that the spreading speed is primarily set by the rate of cytoskeletal protrusion, this mechanism would predict a weaker dependence of the spreading speed on the ligand density. To expose which type of behavior human neutrophils exhibit, we allow the cells to settle onto coverslips pre-coated with different densities of rabbit IgG and analyze their subsequent spreading dynamics. Visualization of the cells with reflection interference contrast microscopy (RICM) allows us to image the cell-substrate contact region with exceptional contrast without the use of labels. In addition to the spreading speed and maximum contact area, this analysis enables us to further characterize the effects of IgG density on phagocytic spreading by quantifying parameters such as the spreading probability, contour roundness, and centroid motion. In an accompanying theoretical study, we leverage these results against computational models of cell-spreading mechanics (Francis and Heinrich, 2022). Overall, we find that, although the strength of adhesion can modulate phagocytic spreading, it is the cellular protrusion that sets the rate of spreading, in support of the protrusive zipper hypothesis.

## Results

### Comparison with bead standards provides density of IgG on glass coverslips

We prepared surfaces displaying the Fc domains of IgG by saturating BSA-coated glass coverslips with anti-BSA antibodies (Fig. 1). To adjust the surface density of human-reactive IgG, we used incubation buffers containing different ratios of polyclonal rabbit IgG and monoclonal mouse IgG-1. The total concentration of IgG was the same in all buffers. Because mouse IgG-1 is hardly recognized by human Fcγ receptors (Lubeck et al., 1985; Shashidharamurthy, 2010; Warmerdam et al., 1990), it served to dilute the amount of human-reactive (rabbit) IgG decorating the coverslips. This approach appeared to produce consistently functionalized substrates for cell-spreading experiments. However, because the rabbit and mouse antibodies may have different affinities for BSA, the ratio of the two types of IgG deposited on the surface is not necessarily the same as in the incubation buffer.

**Fig. 1.**
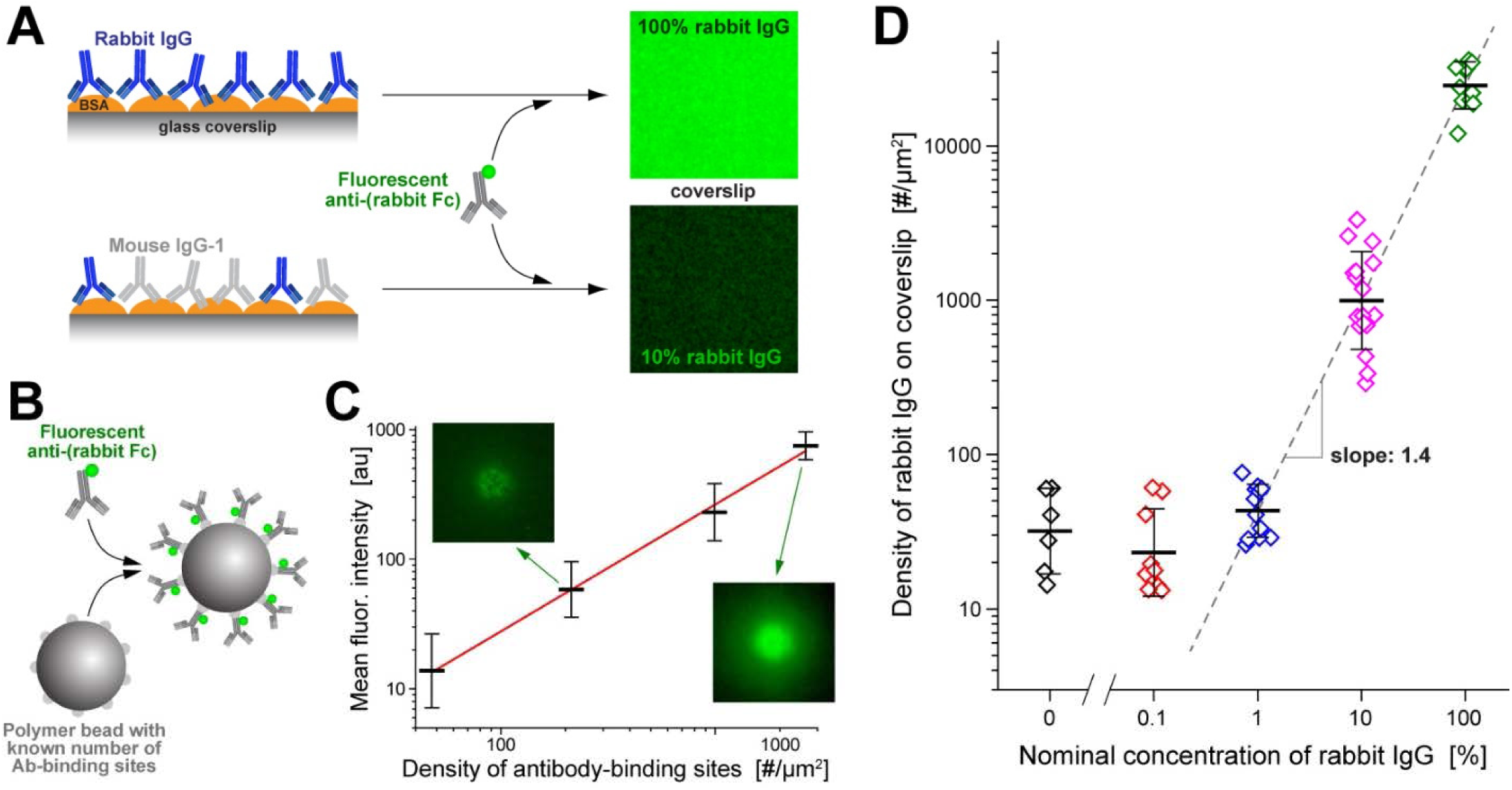
Preparation of substrates with controlled rabbit-IgG surface density. (**A**) BSA-coated glass coverslips that had been incubated with various ratios of rabbit and mouse anti-BSA antibodies were labeled with green-fluorescent secondary antibodies and then imaged on a confocal microscope. (**B**) Standard beads with known numbers of antibody-binding sites were saturated with the same fluorescent secondary antibody as used in (A). (**C**) The fluorescence intensity at the underside of the coated beads was measured with the same confocal-microscope settings as used in (A). The results, along with a straight-line fit (*red line*), are presented as a function of the known density of antibody-binding sites in a log-log plot. Error bars denote the geometric standard deviation. Examples of confocal bead images are included. (**D**) The calibration curve of (C) was used to convert the measured fluorescence intensities of labeled coverslips into the rabbit-IgG density. The geometric mean values (with error bars denoting geometric standard deviation) of the IgG density are plotted as a function of the concentration of rabbit IgG used in the incubation buffer. The result of a linear fit (*dashed line*) to the portion of the data obtained at IgG concentrations of 1% or higher is included.

To quantify the actual amount of deposited rabbit IgG, we labeled IgG-coated coverslips with fluorescent secondary antibody against rabbit IgG (Fig. 1A) and compared the measured fluorescence intensity to a suitable standard. This standard consisted of batches of microspheres pre-functionalized with known numbers of antibody-binding sites (QSC Kit; Bangs Laboratories, Fishers, IN) that we saturated with the same secondary antibody as used to label the coverslips (Fig. 1B). We analyzed *Z*-stacks of confocal images of these beads to determine the intensity of the fluorescent layer at the underside of each bead local to the center of the bead image (Materials and methods; Fig. S1). This analysis provided a calibration curve relating the fluorescence intensity of an essentially flat layer of secondary antibody to the known surface density of rabbit IgG (Fig. 1B). The calibration curve then allowed us to convert measurements of the mean fluorescence intensity of confocal images of labeled coverslips (Fig. 1A) to the density of rabbit IgG deposited on these coverslips. We note that the fluorescence intensity of the most densely coated coverslips lay outside the intensity range of the bead standards; in this case, our analysis involved an extrapolation of the calibration curve.

These measurements confirmed that our method to functionalize coverslips resulted in a reproducible, broad spectrum of controlled rabbit-IgG densities. They also revealed that the relationship between the rabbit-IgG concentration in the incubation buffer—denoted as the “nominal” rabbit IgG concentration throughout this paper—and the actual surface density of rabbit IgG on coverslips was nonlinear (Fig. 1C), indicating that the rabbit and mouse IgGs indeed had different affinities for BSA. Fitting a mathematical model of competitive binding to these data allowed us to estimate that the effective affinity of the used polyclonal rabbit antibody was about 0.4 times that of the monoclonal mouse IgG. The measured surface densities of rabbit IgG ranged from below our detection limit (at 0.1% nominal rabbit-IgG concentration where the surface density was indistinguishable from our negative control) to a maximum density of ∼25,000 rabbit-IgG molecules per μm^2^ (at 100% nominal rabbit-IgG concentration). Assuming a uniform distribution of bound IgG molecules, the maximum surface density corresponded to an area of roughly 40 nm^2^ occupied by each rabbit IgG, consistent with a dense, essentially contiguous layer of IgG coating the coverslips.

### ells are more likely to spread on higher densities of IgG

We used an IgG-coated coverslip as the bottom of our assembled experiment chamber (Fig. 2A,B). A plastic insert with funnel-shaped vertical thru holes served as the chamber’s top (Fig. 2A). The chamber was filled with Hanks’ balanced salt solution supplemented with 2% human serum albumin and placed on an inverted research microscope set up for epi-illumination. This design allowed us to introduce small volumes of a suspension of isolated human neutrophils at non-overlapping locations through the chamber ceiling. Each deposited cell population dispersed to some extent by diffusion while sinking to the chamber bottom. The cells settled onto the functionalized coverslip within ∼5 minutes, usually without piling into clusters.

**Fig. 2.**
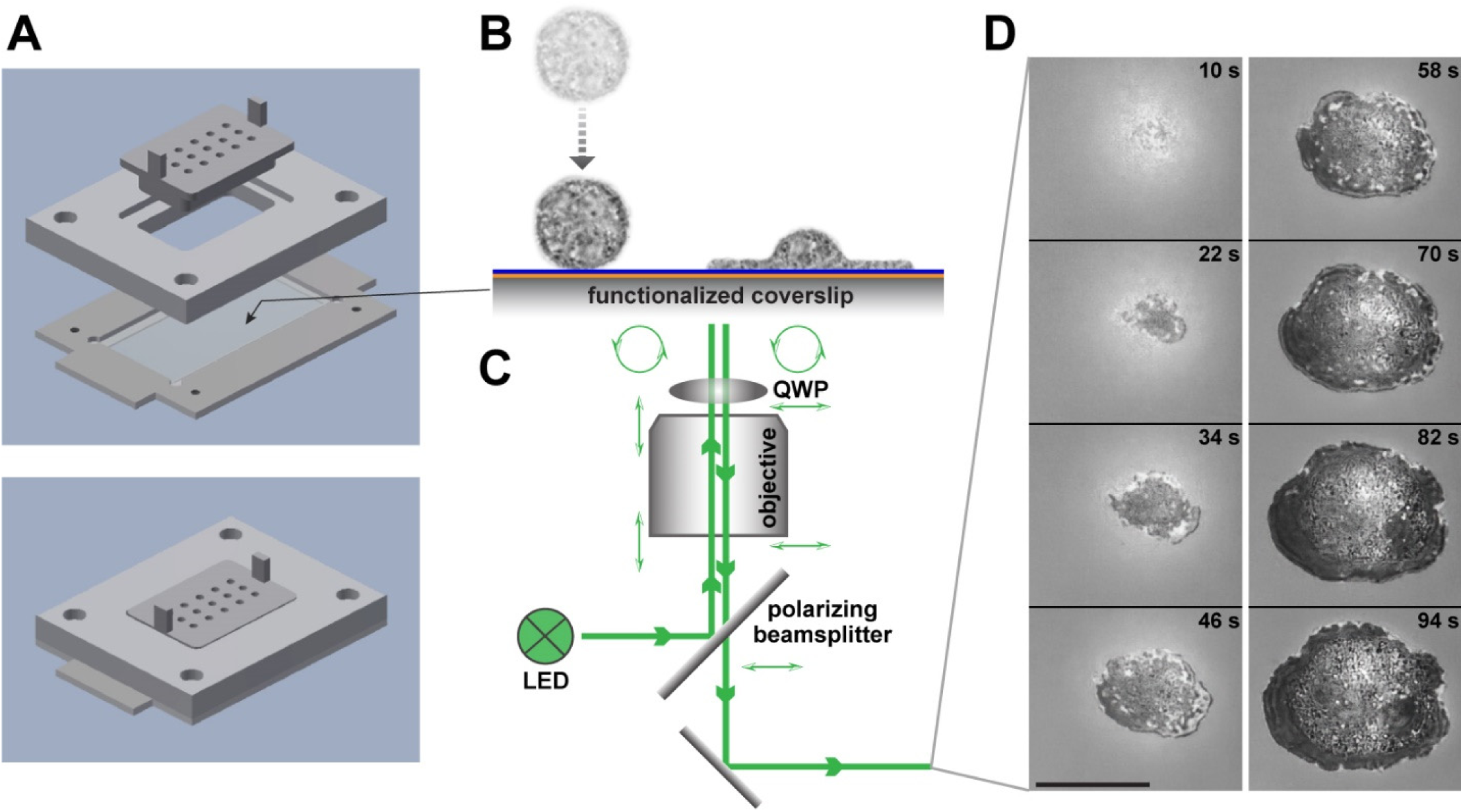
Illustration of a frustrated phagocytic spreading experiment. (**A**) Parts and assembly of the experiment chamber. (**B**) After settling onto the functionalized coverslip, initially passive, round cells recognize the deposited antibodies and spread along the surface. (**C**) Reflection interference contrast microscopy allows us to visualize the cell-substrate contact area label-free at high resolution. The combination of polarizing beamsplitter and quarter-wave plate (QWP) ensures that the light components reflected off the coverslip top surface and the cell are the dominant contributions to the recorded interference pattern. (**D**) Example snapshots from a series of recorded video images of the dark cell-substrate contact area (see also Movie S1). Timestamps are included. The scale bar denotes 10 μm.

We imaged the cells using reflection interference contrast microscopy (RICM, Fig. 2C,D, Movie S1). The interference patterns produced by RICM markedly enhanced the visibility of contact regions between the primary light-reflecting interface—the top surface of the coverslip— and the underside of reflective objects resting on this surface. Regions of direct cell-substrate contact generally appeared as clearly outlined, distinctive dark patches. It is important to bear in mind though that light-reflecting structures inside a cell, such as granules, can produce brighter spots within the dark contact region that could be misinterpreted as areas where part of the cell has detached from the substrate. Close inspection of the interference patterns revealed that in our experiments, contiguous contact regions did not appear to contain local areas of cell-substrate separation.

Contact with the substrate caused a fraction of the cells to start spreading either immediately or after some lag time. Despite the mixed response of the cells in a given population—reflecting cell-to-cell variation typical for the behavior of primary human immune cells—we found that the overall proportion of spreading to non-spreading cells strongly depended on the density of rabbit IgG coating the coverslip. We quantified the cells’ commitment to spreading on a given surface in terms of the “spreading probability”, i.e., the ratio between the number of spreading cells and the total number of observed cells. We consistently evaluated this ratio 30 minutes after depositing the cells, defining “spreading cells” as those cells whose cell-substrate contact area had reached at least 100 µm^2^ at this time.

The measured spreading probabilities ranged from 0.16 on the lowest density of surface-bound rabbit IgG to about 0.85 on the highest density (Fig. 3). The result for the lowest density was not significantly different from the spreading probability on substrates presenting pure mouse IgG-1 (negative control). While this result may be attributed in part to spurious recognition of mouse IgG-1 by human neutrophils, it also highlights that neutrophil populations almost always contain cells that can be activated by non-specific interactions with otherwise non-stimulatory target surfaces. On the other hand, almost all cells spread on substrates coated with high-density, 100% rabbit IgG, confirming that antibody-coated targets generally draw a vigorous response by human neutrophils.

**Fig. 3.**
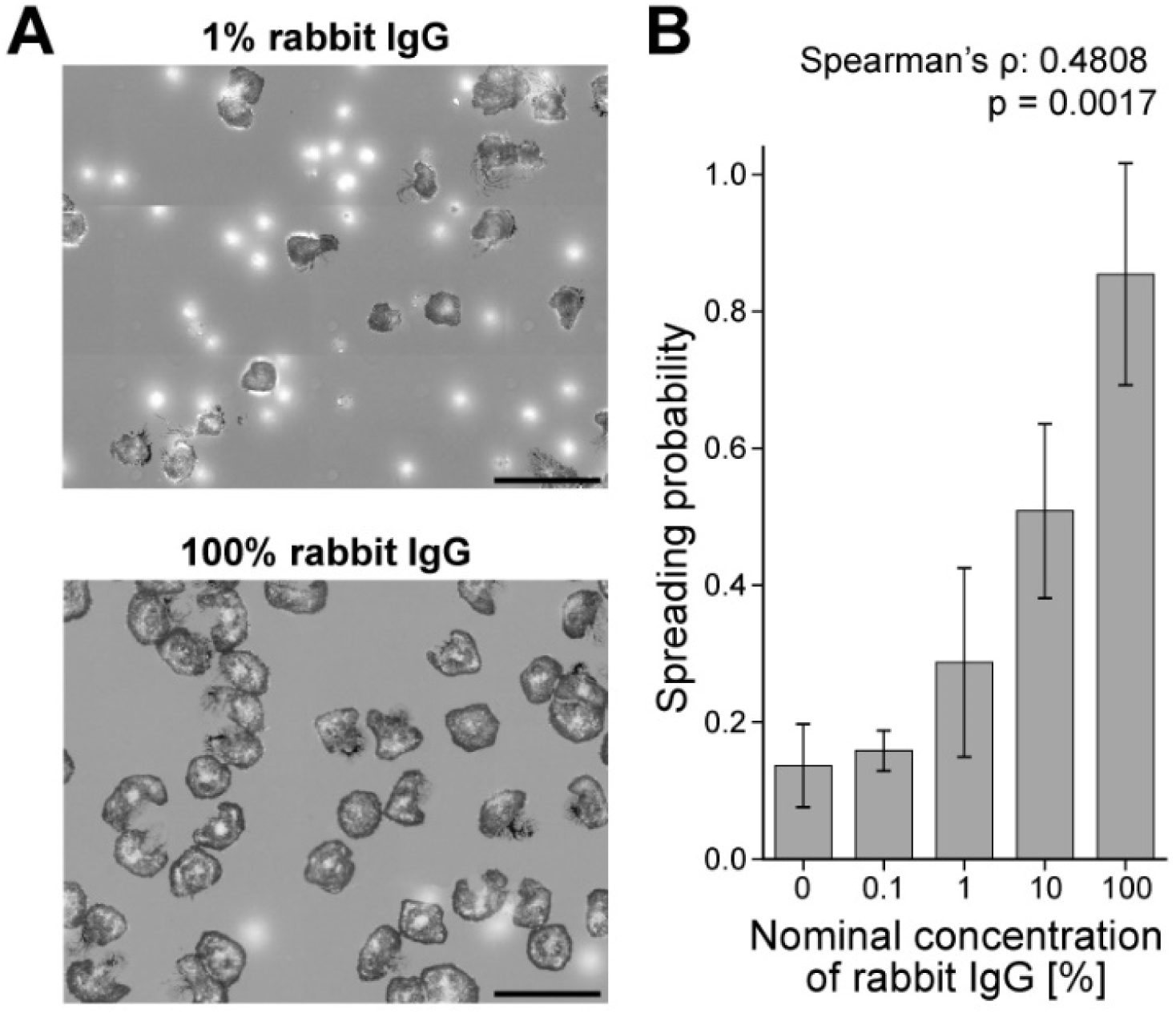
Dependence of the spreading probability on the surface density of IgG. (**A**) At low surface densities of rabbit IgG, the majority of deposited cells do not spread and appear as bright, out-of-focus spots in our RICM images (*top panel*). In contrast, almost all cells spread on surfaces coated with the highest density of rabbit IgG (*bottom panel*). Scale bars denote 50 μm. (**B**) The spreading probability depends strongly on the nominal concentration of rabbit IgG, confirming that the observed cell response is IgG-specific. Spearman’s rank correlation coefficient ρ is reported with the associated *p*-value for the null hypothesis ρ = 0.

We limit our remaining analysis to spreading neutrophils. Thus, when interpreting the following results, it is important to bear in mind that they were obtained with selected subpopulations of cells, and that the fraction of cells making up each subpopulation depended on the cell response to a particular surface density of rabbit IgG.

### The cell-substrate contact region of spreading cells is essentially radially symmetric

After settling on the bottom coverslip, individual neutrophils typically formed one or two initial attachment spots with the IgG-coated substrate. In cases were more than one spot was visible, the growing spots quickly merged into a single, contiguous region of cell-substrate contact. Subsequently, the contact region of spreading neutrophils generally had a roughly circular shape that expanded in a radially symmetric manner, as expected for phagocytic spreading (Fig. 4). It typically took neutrophils only about 2 minutes from the start of spreading to form contact footprints with diameters as large as 20 μm.

**Fig. 4.**
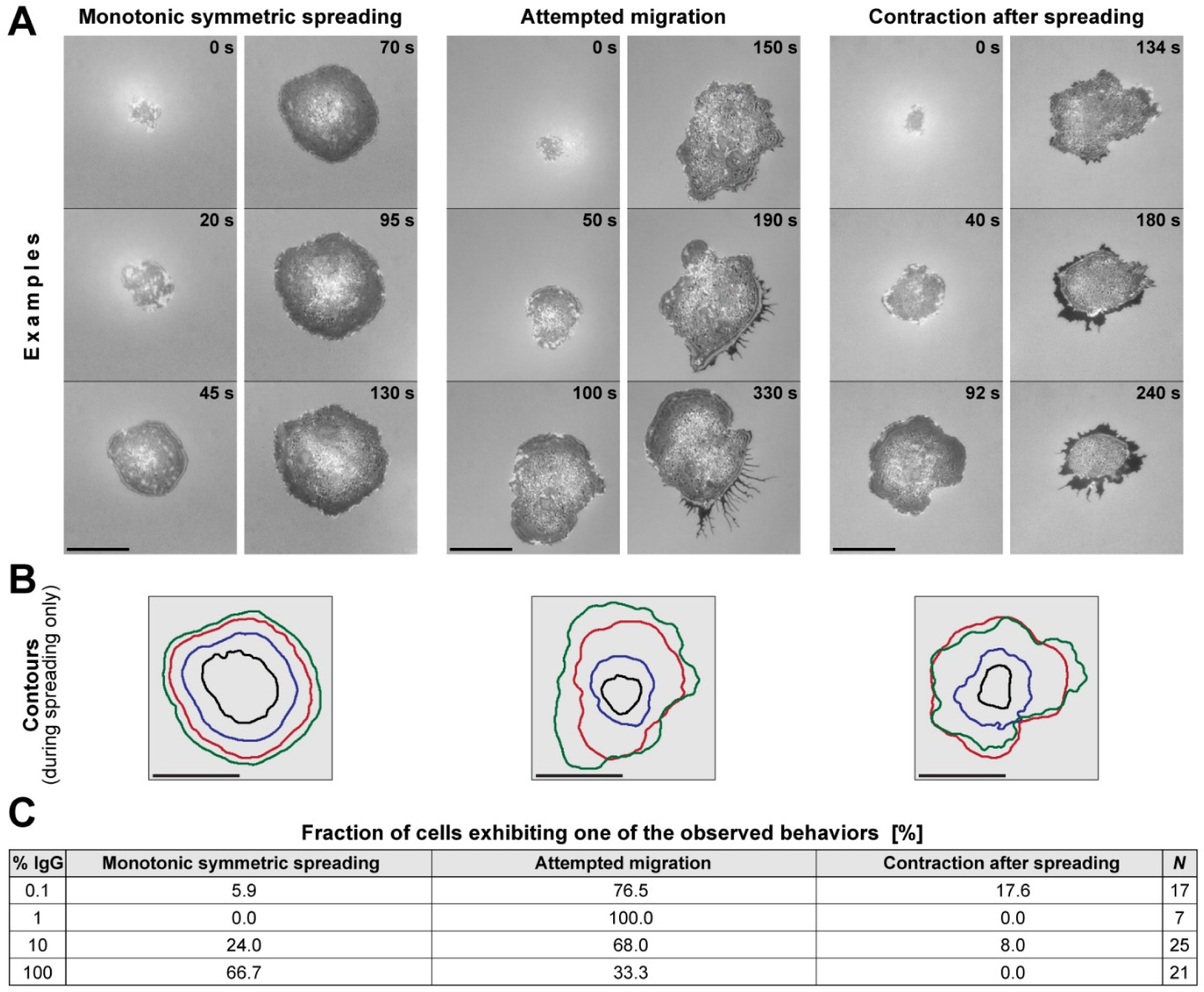
Overview of the human neutrophil response to IgG-coated surfaces. (**A**) The first four snapshots of each video sequence illustrate the common morphology of the cell footprint observed during the outward-spreading phase of almost all cells (see also Movie S1). In contrast, three qualitatively distinct types of post-spreading cell behavior were observed, as illustrated in the last two snapshots of each video sequence. Time stamps are included. Scale bars denote 10 μm. (**B**) Contours of the cell footprint of the first four images of the respective video sequences of (A) demonstrate the roughly symmetric cell morphology during spreading. Scale bars denote 10 μm. (**C**) The table reports the fractions of cells that exhibit one of the three types of post-spreading behavior for each of the tested IgG concentrations. On low densities of IgG, the majority of cells attempted to migrate after reaching a maximum cell-substrate contact area. In contrast, on high IgG densities the majority of cells remained in place and exhibited little further shape changes.

After the contact area reached an apparent plateau, neutrophils exhibited a spectrum of different behaviors. Some remained in place but continued to modify the cell-substrate contact region, gradually expanding or contracting it, or altering its shape while leaving its net area essentially unchanged. Other cells appeared to adopt a more migratory phenotype, attempting to crawl away from the original location of spreading. Common to most post-spreading behaviors was the formation of trailing membrane extrusions at retracting cell regions. These structures connected the main cell body to focal cell-substrate adhesion sites from which the moving cell often was unable to detach (Fig. 4). We have not attempted to analyze this highly variable post-spreading behavior beyond a coarse empirical classification (Fig. 4), demonstrating that cells spreading on higher densities of IgG were less likely to try to leave their original spot of spreading (Fig. 4C).

### Density of IgG does not affect spreading speed but weakly correlates with extent of spreading

Our quantitative comparison of neutrophils spreading on different densities of IgG is based on the analysis of image sequences recorded at intervals of 2 s. We semi-automatically traced the outline of the cell footprint in each image and stored the resulting polygons of cell peripheries (Fig. 5A). Our primary quantity of interest was the area circumscribed by these polygons, i.e., the cell-substrate contact area. Plots of the contact area as a function of time revealed a comparatively slow, brief initial spreading phase, followed by a roughly linear expansion of the contact footprint, and eventually, a gradual approach to a plateau of maximum contact area (Fig. 5A-C).

**Fig. 5.**
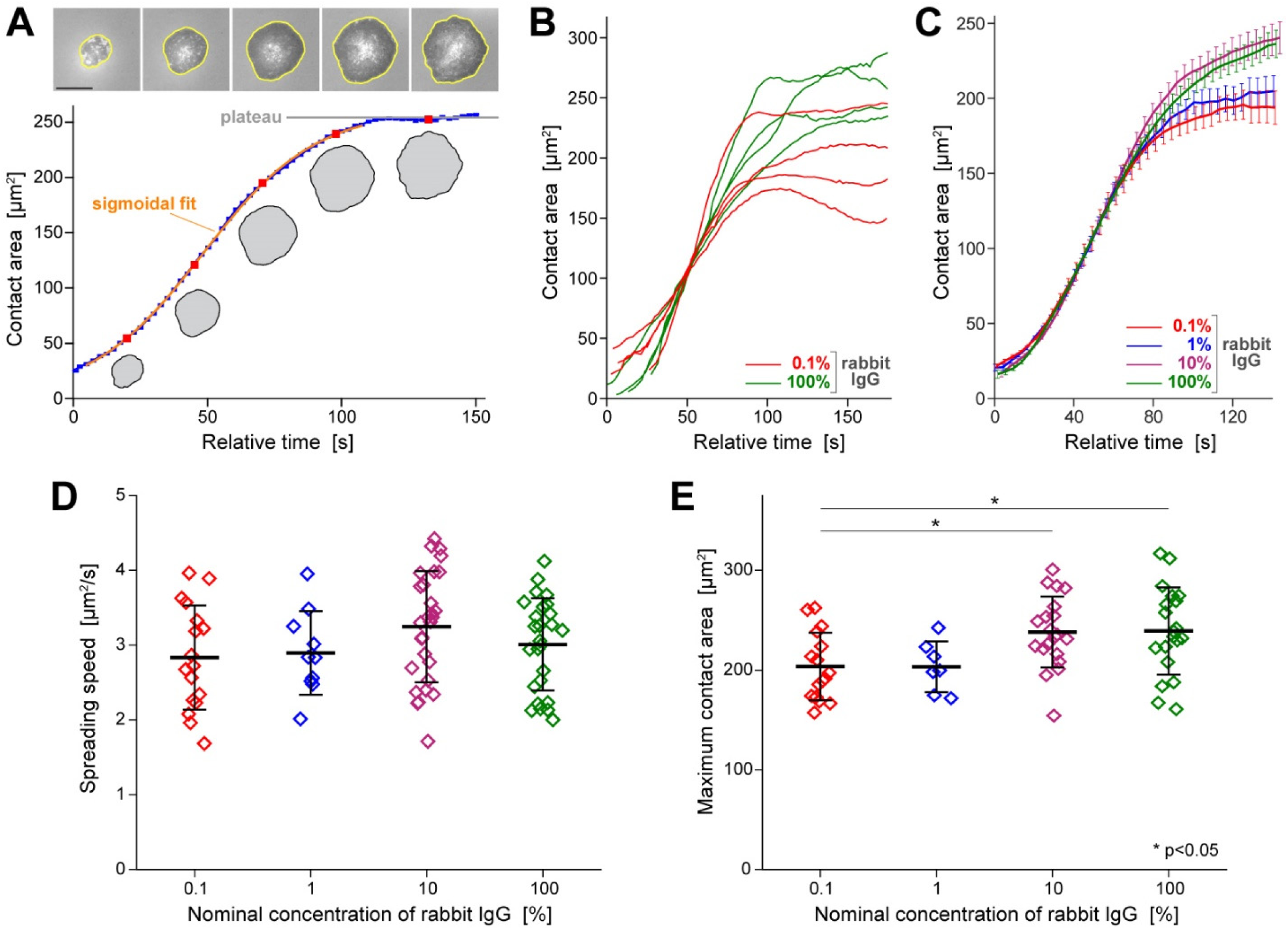
Quantitative analysis of the cell-substrate contact area of spreading cells. (**A**) The contact area is determined from a polygonal trace of the circumference of each cell footprint (*yellow polygons* in the examples included at the top; scale bar denotes 10 μm). A suitable fit to the time-dependent contact area yields the (maximum) spreading speed. The plateau value of the contact-area curves provides the maximum contact area. (**B**) Representative examples of contact-area-versus-time curves measured on the lowest (*red*) and highest (*green*) IgG densities illustrate the variability of the dynamical behavior of the cell footprint. (**C**) Average curves of all suitable area-versus-time measurements obtained for each tested rabbit-IgG density expose a largely conserved spreading speed. On the other hand, the maximum contact area appears to reach greater values on high IgG densities. Error bars denote standard errors. (**D**) The summary of all spreading speeds measured over three orders of magnitude of the IgG concentration confirms that the spreading speed is independent of the surface density of IgG. (**E**) The summary of maximum contact areas reveals a small but significant increase of the contact area at high densities of IgG. Error bars in (D) and (E) denote standard deviation.

We define the speed of spreading as the value of the steepest slope of a given contact-area-versus-time graph. We used sigmoidal or linear fits to such curves to determine the spreading speeds in suitable experiments. Remarkably, the average spreading speeds, measured over a 1000-fold range of nominal IgG concentration, were all close to 3 μm^2^/s, exhibiting no significant differences on different IgG densities (Fig. 5D).

The similarity in neutrophil behavior during the rapid-spreading phase also was evident in example curves measured for cells spreading on 0.1% versus 100% rabbit IgG (Fig. 5B). On the other hand, the collection of graphs exposed an apparent difference between the maximum contact areas occupied by cells spreading on these two IgG densities. Most cells spreading on 100% IgG reached contact areas in the range of 200-300 μm^2^, whereas on the substrates coated with the lowest IgG density (0.1%), the maximum contact area tended to lie in the range of 150-250 μm^2^. The statistical analysis of all measurements confirmed that the mean maximum contact areas on surfaces coated with 10% or 100% IgG were indeed significantly larger than those on 0.1% IgG (Fig. 5E).

Recalling that originally spherical cells (with diameter *D*_0_) must increase their apparent surface area when spreading on a substrate, it is instructive to convert the measured area of cell-substrate contact footprints into estimates of the overall cell-surface area (*A*) required to accommodate such deformations. Our conversion assumes that the cell volume remains constant during spreading, and that the cell-substrate contact region is circular, denoting its diameter by *D*_*c*_. Furthermore, we approximate the shape of the upper, free surface of the spread cell as a spherical cap, which is the geometry with the smallest possible surface area for the given volume and a circular contact footprint. For this simplified geometry, one can calculate the ratio between the final area *A* and the initial cell-surface area *A*_0_=π*D*_0_^2^ using the following prescription

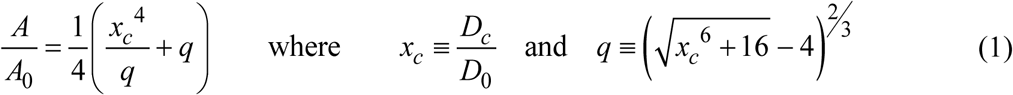

The diameter of most non-adherent, resting human neutrophils lies in the range of 8.5-9 µm. With the choice of *D*_0_ = 8.75 µm, Eq. (1) predicts that a typical neutrophil will need to expand its surface area to ∼210% of its initial area in order to form a contact footprint whose diameter is twice as large as the cell’s resting diameter (where *x*_*c*_ = 2 and *D*_*c*_ = 17.5 µm).

The average of the 10 largest cell-substrate contact areas measured in our experiments is 359 μm^2^, corresponding to a typical footprint diameter of *D*_c_ = 21.4 µm. Thus, we estimate that the most-spread neutrophils have expanded their apparent surface area to about 303% of their resting area during our frustrated-phagocytosis experiments, in agreement with previous findings (Hallett and Dewitt, 2007; Herant et al., 2005; Lee et al., 2015; Simon and Schmid-Schonbein, 1988).

### Higher IgG density results in more uniform, concentric spreading

The results of the previous section provide compelling support for the protrusive-zipper hypothesis. A detailed analysis of additional features of the cell behavior is likely to produce further insight into the fundamental mechanisms of phagocytic spreading. We next examined how the IgG density affects the roundness and type of motion of the cell footprint during spreading.

We assessed the roundness of the cell-substrate contact region in terms of the ratio of the radii of two circles defined by the outline of this region, i.e., the largest inscribed circle and the smallest circumscribed circle (Fig. 6A). The value of this roundness measure equals 1 for circular contours and is smaller otherwise, with lower values corresponding to less round contours. For each spreading cell, we measured the roundness of individual contours within the phase of fastest contact-area growth (where the spreading speed was greater than 0.4 µm^2^/s), then applied a moving average filter with a window size of 8 s and finally determined the maximum of these averages. This approach yielded roundness values in the range of roughly 0.6-0.9 that exhibited no significant differences between surfaces coated with different densities of IgG (Fig. 6B). Thus, the cell contours generally were characterized by a moderate roundness that was independent of the IgG density, indicating that phagocytic spreading proceeded in a more or less uniform fashion on all tested surfaces.

**Fig. 6.**
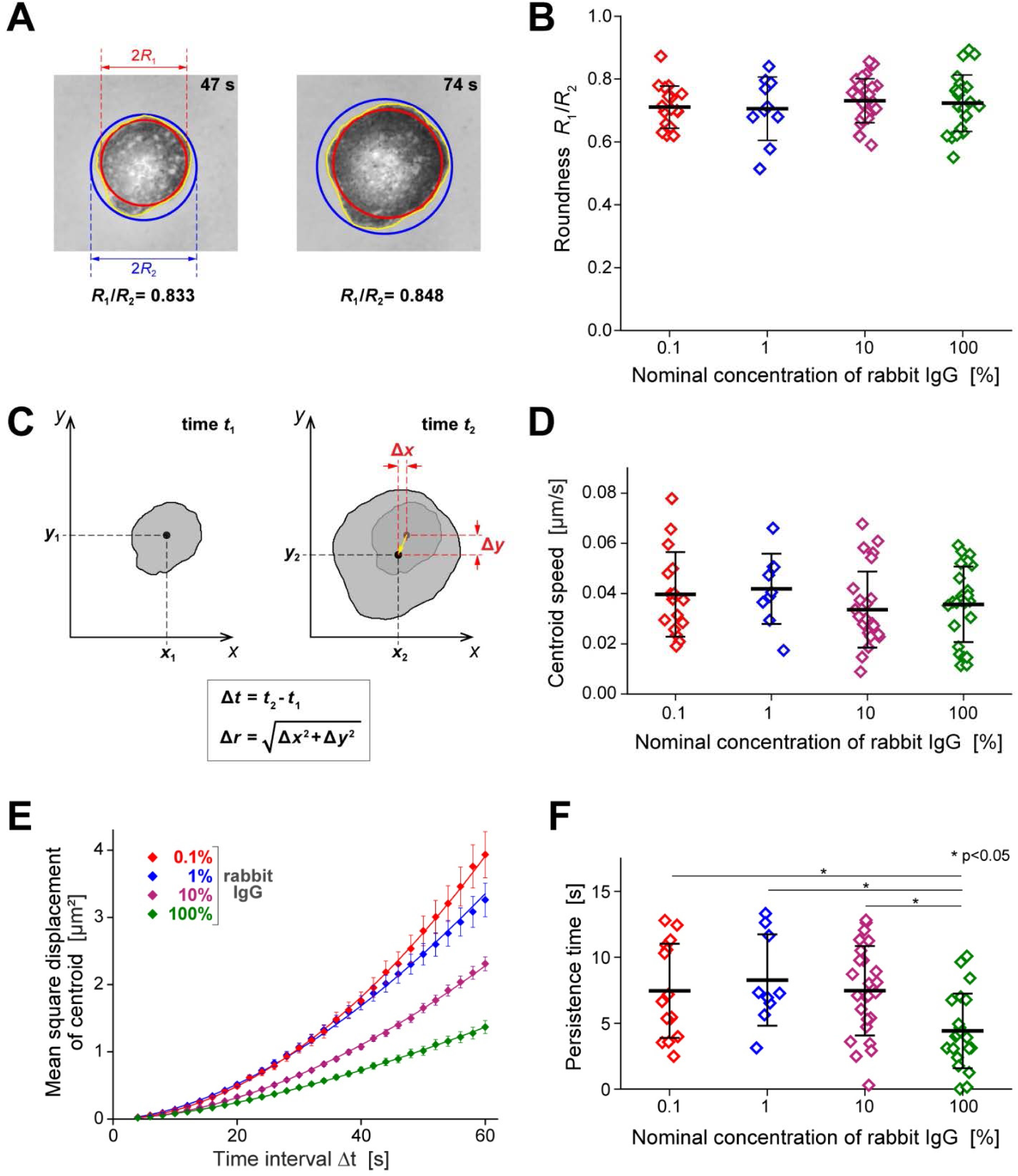
Roundness and type of motion of the footprints of spreading cells. (**A**) The roundness of the cell-substrate contact region was assessed in terms of the ratio between the radii of the largest inscribed (*red*) and the smallest circumscribed (*blue*) circles, as illustrated with two examples. (**B**) The summary of all roundness measurements reveals no significant differences on surfaces coated with different IgG densities. (**C**) Illustration of our measurement of the displacement Δ*r* of the centroid of the cell footprint during the time interval Δ*t*. (**D**) The summary of all centroid-speed measurements reveals no significant differences on surfaces coated with different IgG densities. (**E**) The mean square displacements of the centroid as a function of Δ*t* (averaged over all suitable measurements obtained for each tested rabbit-IgG density; error bars denote standard errors) allow us to quantify the degree of randomness of the centroid motion. The included solid lines show power-law fits of the form *y*°=°*ax*^*b*^ to the data. (**F**) The summary of all measured directional persistence times reveals that on the highest density of IgG, the cell footprint tended to move in roughly the same direction for significantly shorter times than on lower IgG densities. For more details see the text.

The roundness measure does not capture unambiguous information about possible lateral displacements of the cell footprint. To quantify the latter, we analyzed the positions of the centroid of the cell-substrate contact region of spreading cells in suitable video images. We defined the centroid displacement Δ*r* as the Euclidean distance by which the centroid position moved during a given time interval Δ*t* (Fig. 6C). We used these measurements to estimate the instantaneous velocity of the centroid motion in terms of the ratio Δ*r*/Δ*t* evaluated for successively recorded centroid positions. For each spreading cell, we then averaged the instantaneous values over the active spreading phase. Most of the resulting average centroid speeds were in the range of 0.01-0.06 µm/s and exhibited no significant differences between substrates coated with different IgG densities (Fig. 6D).

In addition to evaluating the centroid speed, we also used the measured centroid positions to assess the type of motion of the cell footprint. Now choosing Δ*t*-values in the range from 4 s to 60 s, we averaged the squared centroid displacements of a given cell over all pairs of video frames whose recording times differed by Δ*t*. We then calculated the averages of these mean square displacements as a function of Δ*t* for the whole population of cells spreading on a given substrate (Fig. 6E). Power-law fits of the form *y*°=°*ax*^*b*^ to plots of these mean square displacements as a function of Δ*t* allow for a rough classification of the type of centroid motion. An exponent of *b*°=°1 of such a fit implies a random motion with Gaussian-distributed individual displacements, as is typical, for example, for diffusion. If the exponent approaches 2, the motion generally is interpreted as more migratory. The exponent of the power-law fit to the measured mean square centroid displacements was 1.57 on the highest density of IgG, whereas on the lowest IgG density it was 1.86, suggesting that lateral displacements of cells spreading on the highest IgG density were more random, i.e., the cells were less likely to transiently explore the substrate in a preferred direction.

The lack of a clear correlation between IgG density and centroid speed (Fig. 6D) contrasted with the observed differences in centroid displacement (Fig. 6E), suggesting that on some substrates, moving cells changed direction more frequently than on others. We verified this assumption by assessing the directional persistence of centroid displacements in terms of the decay of the time-dependent autocorrelation function of the tangents of the centroid trajectory (Gorelik and Gautreau, 2014). The resulting “directional persistence time” (Materials and methods) exposes for how long a cell footprint continues to move in roughly the same direction. The measured persistence times generally were small and exhibited a large spread; however, they were significantly shorter (∼4.4 s on average) on coverslips coated with the highest density of IgG than on lower IgG densities (∼7.5-8.3 s) (Fig. 6F). The averages of the closely related “directional persistence distances” (product of centroid speed and persistence time) traversed in a preferred direction were small, falling in the overall range of 0.16-0.36 µm, and exhibiting a significant difference between substrates coated with nominal IgG concentrations of 100% and 1% (data not shown). In summary, transient lateral polarization and directional movement of the cells, while generally unlikely, indeed met with stronger resistance on higher densities of IgG.

This analysis confirmed that the observed type of spreading motion was specific to the phagocytic ligand IgG. Overall, cells spreading on IgG tended to expand the contact region in a uniform, concentric manner without extending substantial exploratory protrusions in preferred directions. As shown earlier (Fig. 4), this behavior may change once the area of the cell-substrate contact region has reached an apparent plateau.

### Computer simulations reinforce mechanistic insights gained from experiments

The comparison of experimental results with theoretical predictions is an essential approach to test mechanistic hypotheses and decide whether observed correlations are coincidental or causative. In a companion study, we have translated different hypotheses about the driving force of phagocytic spreading into mathematical models and examined the mechanical ingredients that such models require to reproduce the cellular behavior (Francis and Heinrich, 2022). This section provides a broadly accessible summary of our comparison between the measured dynamics of the growth of the cell-substrate contact area and theoretical predictions based on the Brownian zipper and the protrusive zipper models, respectively.

Details of our mathematical models are provided in the companion paper (Francis and Heinrich, 2022). In short, we consider an immune cell as an axisymmetric body with uniform surface tension that is filled with a highly viscous, incompressible fluid (Tran-Son-Tay et al., 1991; Yeung and Evans, 1989). The model incorporates adhesive interactions between the cell surface and a flat, rigid substrate in terms of a short-range attractive force that acts to pull the cell onto the substrate in an irreversible manner. This basic version of the model captures the essential assumptions of the Brownian zipper hypothesis.

To introduce active cellular protrusion into this framework, we postulate that fresh contact between the substrate and the cell surface results in a transient, outward-pushing force local to the leading edge of the cell-substrate contact region. We do not attempt to model the microscopic origin of this protrusive force, which would have to account for receptor-induced signaling, spatiotemporal redistribution of messenger molecules, translation of biochemical cues into mechanical stresses, etc. Instead, our semi-empirical approach lumps these highly complex processes into the rationale that receptor activation due to fresh cell-substrate contact ultimately causes the protrusive force. We also assume that the magnitude of protrusion decays as a function of the time elapsed after the most recent receptor-ligand binding event. One objective of our theoretical study is to establish what distribution and magnitude of mechanical stresses can account for the experimental results, and whether the predicted stresses are physically realistic and biologically plausible. The model version that includes the protrusive force represents the protrusive zipper hypothesis.

As reasoned earlier, adhesion-driven, passive spreading, which forms the core of the Brownian zipper hypothesis, should result in a stronger correlation between the IgG density and the spreading speed than active, primarily protrusion-driven spreading. To verify this common-sense expectation and place it on sound quantitative grounds, we simulated the dependence of the spreading speed on the IgG density using both versions of our mathematical model (Fig. 7). The simulations indeed revealed a dramatic difference in predicted cell-spreading behavior between these two models. For realistically chosen parameter values, the Brownian zipper model predicts a strong dependence of the cell spreading speed on the density of IgG binding sites, and the spreading speeds on low IgG densities are predicted to be much slower than observed experimentally. In contrast, the spreading speed predicted by the protrusive zipper model hardly changes over a 1000-fold range of nominal IgG concentrations and is similar to experimental results. Comparison of the predictions of Fig. 7D with the measurements shown in Fig. 5C,D clearly supports the conclusion that the protrusive zipper hypothesis successfully captures the basic mechanisms underlying phagocytic spreading, whereas the Brownian zipper hypothesis does not.

**Fig. 7.**
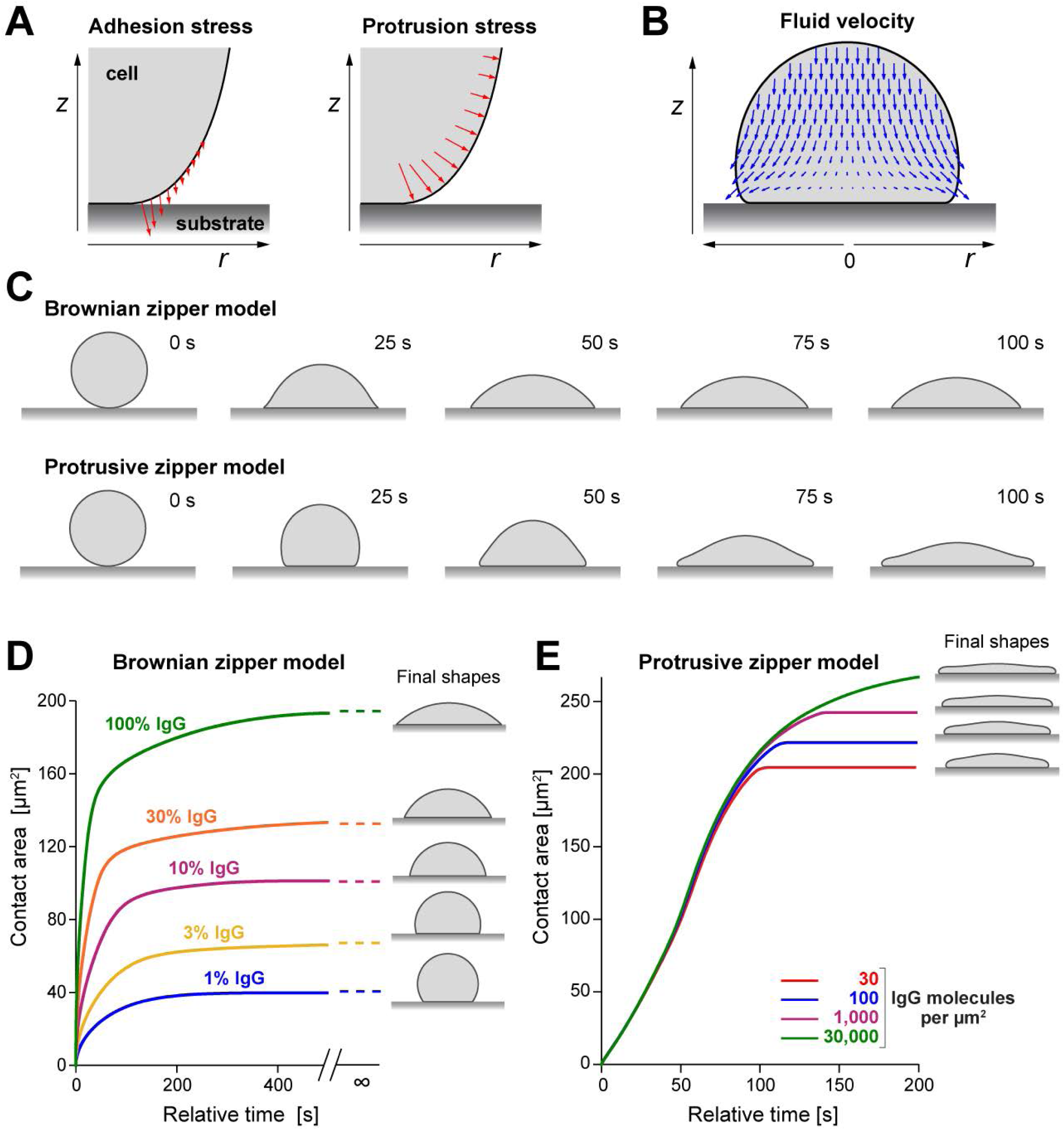
Summary of pertinent predictions by two different mechanistic models of cell spreading. (**A**) The models incorporate attractive interactions between the cell surface and the substrate via short-range adhesion stress (*left panel*). The protrusive-zipper model postulates that the cell interior generates protrusion stress local to the leading edge of the spreading cell in a manner that depends on fresh contact between the cell and substrate (*right panel*). (**B**) A snapshot of a computer simulation of a spreading model cell illustrates the motion of the viscous cell interior according to the Stokes equations for creeping flow. (**C**) Time series of cross-sectional shapes of spreading model cells illustrate morphological differences between the predictions of the two considered models. (**D**) The Brownian-zipper model predicts that both the cell-spreading speed as well as the maximum contact area exhibit a strong dependence on the surface density of IgG coating the substrate. The final shapes are predicted to be spherical caps in this case. These predictions fail to reproduce experimental observations such as presented in Fig. 5. (**E**) In agreement with our experimental results (Fig. 5C,D), the spreading speeds predicted by the protrusive-zipper model are essentially independent of the IgG density, and the IgG dependence of the maximum contact area is much weaker than in (D).

The dependences of the maximum contact area on the IgG density (Fig. 7D) reveal another difference between the two model versions. Recall that our experiments exposed a significant increase of the maximum contact area at higher IgG densities (Fig. 5E). However, this increase is much smaller than the large differences between the plateau values of the graphs predicted by the Brownian zipper model (Fig. 7D). On the other hand, the predictions of the protrusive zipper model again provide a notably better match with our experimental observations (Fig. 7D).

## Discussion

The rapid execution of complex cellular motions is one of the most amazing feats of immune cells, and critical to phagocytosis, cell migration, and other vital processes. However, precise descriptions of such motions and, more importantly, understanding of the underlying mechanisms, remain scarce. The quantitative analysis of frustrated phagocytosis presented in this paper is part of a larger research area that focuses on the biomechanics of cells spreading on surfaces, comprising both experimental work (Dobereiner et al., 2004; Dubin-Thaler et al., 2004; Dubin-Thaler et al., 2008; Hategan et al., 2004; Janmey et al., 2021; Reinhart-King et al., 2005; Wolfenson et al., 2014) as well as theoretical studies (Chamaraux et al., 2005; Cuvelier et al., 2007; DiNapoli et al., 2021; Etienne and Duperray, 2011; Fardin et al., 2010; Frisch and Thoumine, 2002; Odenthal et al., 2013; Xiong et al., 2010). Our integrative experimental/theoretical approach expands most previous analyses by quantifying how the ligand density affects the cells’ spreading behavior, and to what extent adhesive versus protrusive forces contribute to the cell motion. In conjunction with our theoretical companion paper, this approach has allowed us to resolve controversial mechanistic notions about the spreading behavior of immune cells.

Our experimental strategy has been to expose human neutrophils to target surfaces that presented a range spanning three orders of magnitude in density of deposited IgG. The strong dependence of the cells’ spreading probability on the IgG density confirms that the ligation of Fcγ receptors is the original cause of spreading. It also suggests that a threshold in cell stimulation needs to be reached to induce spreading. The most likely type of stimulating trigger appears to be a minimum number of engaged receptors. Alternatively, the primary stimulus could be the spacing between engaged receptors, in which case their total number might play a lesser role. Our current setup does not allow us to distinguish between these two alternatives. It also is important to bear in mind that the threshold level that triggers spreading can vary considerably from cell to cell, suggesting that it is affected by other factors such as the degree of quiescence of individual neutrophils.

The absence of a significant correlation between the IgG surface density and the spreading speed of the cells is perhaps the most consequential result of our experiments. It provides a counterexample to the notion that spreading might be driven by strong cell-substrate adhesion, thus discrediting the Brownian zipper hypothesis in the case of neutrophils and similar cells. In contrast, the protrusive zipper hypothesis intuitively provides a better explanation of our results. The strongest support for the validity of the latter hypothesis comes from the agreement between computer simulations based on the protrusive zipper model and our experiments performed on different substrates. Our finding agrees well with reports that during isotropic fibroblast spreading, the speed of spreading depended on protrusive stress generated by the actin cytoskeleton (Dubin-Thaler et al., 2008; Fardin et al., 2010; Xiong et al., 2010).

It is worth noting that the spreading speed of macrophage-like cells of the J774 murine cell line on IgG-coated surfaces also was found to be largely independent of the IgG density (Kovari et al., 2016). Thus, our conclusion that cell spreading is primarily driven by an active protrusive force likely holds for motile immune cells in general. It also agrees with a previously reported “all-or-nothing” signaling response during IgG-mediated phagocytosis by macrophages (Zhang et al., 2010), as well as with other studies in which macrophages readily consumed particles coated with low densities of IgG (Ben M’Barek et al., 2015; Pacheco et al., 2013). On the other hand, it conflicts with computer simulations that neglected the role of cytoskeleton-driven protrusion and predicted that phagocytosis should proceed most quickly at an optimal ligand density (Richards and Endres, 2014).

Our finding that the spreading probability, but not the spreading speed, depends on the IgG density reveals that beyond the stimulus threshold that causes a cell to commit to spreading, further strengthening of the stimulus appears to have little effect on the mechanical cell response. In other words, the spreading speed becomes decoupled from the stimulus strength at some point. This raises the interesting question at which step in the sequence of processes leading from receptor ligation to cell movement the decoupling occurs. Does receptor-induced signaling reach a limiting level beyond which the activation of additional receptors is ignored? Is there a bottleneck built into the redistribution of messenger molecules? Or are the cell’s resources to generate additional protrusive force exhausted? Answers to questions like these, although beyond the scope of this study, are an essential part of a comprehensive understanding of vital immune-cell functions.

Our analyses of the geometry and type of motion of the contact footprint, as well as the post-spreading behavior of neutrophils, round out this study, allowing us to paint a detailed picture of the events following cell contact with an IgG-coated surface. After reaching the surface, an initially quiescent cell may undergo detectable Brownian motion and/or be dragged along the surface by convection until cell-substrate adhesion causes its arrest. Cell arrest is usually faster—often immediate—on surfaces coated with higher densities of IgG, suggesting that it is mediated by specific bonds between immobilized Fc domains and Fcγ receptors of the cell. It is worth bearing in mind though that neutrophils can arrest even on negative control surfaces through nonspecific interactions.

Ligation of Fcγ receptors ultimately leads to cell activation and spreading. Many intermediate mechanistic details of this response remain uncertain, but our results strongly suggest that an active protrusive force generated by the cell local to the periphery of the contact footprint is the main driver of spreading. There is ample qualitative evidence supporting the cytoskeleton’s key role in the generation of this force through local actin polymerization and cross-linking, both generally (Footer et al., 2007; Kovar and Pollard, 2004; Parekh et al., 2005) as well as in the specific context of phagocytosis (Allen and Aderem, 1996; Freeman and Grinstein, 2014; Jaumouille et al., 2019; Ostrowski et al., 2019). For example, F-actin accumulation at the front of the phagocytic cup has been shown to be correlated with force production during macrophage phagocytosis (Nelsen et al., 2020). Novel techniques to measure phagocytic forces with high resolution hold great promise for illuminating the detailed mechanisms linking actin remodeling to force generation (Vorselen et al., 2021; Vorselen et al., 2020). It seems plausible to ascribe the local character of this cytoskeletal remodeling to localized signaling triggered by fresh cell-substrate contact.

Although cell-substrate adhesion is not the primary cause of spreading, it still plays a critical role by securing and stabilizing fresh contact regions, thus aiding further cell motion parallel to the surface. A subtle detail in the cell’s control of its motion is the coordination of structural linkages between the actin cytoskeleton and the plasma membrane. Weakening of such linkages at the protruding front is likely to facilitate local forward displacements of the cell surface. A molecular example of such behavior is the calcium-dependent disruption of FERM-mediated linkages by calpain (Dewitt and Hallett, 2002; Dewitt and Hallett, 2020; Roberts and Hallett, 2019). In contrast, in regions of established cell-substrate contact, strengthening of membrane-cytoskeleton linkages prevents the main cell body from being lifted away from the surface, while also providing bracing support to the protrusive motion along the surface (Barger et al., 2019; Herant et al., 2011; Jaumouille et al., 2019; Jaumouille and Waterman, 2020). Several molecular players that either link integrins to the cytoskeleton (e.g., talin, paxillin) or directly link the membrane to the cytoskeleton (e.g., myosin I, ezrin) have recently been shown to colocalize with the neutrophil or macrophage phagocytic cup and facilitate directional force generation (Barger et al., 2019; Jaumouille et al., 2019; Ostrowski et al., 2019; Roberts et al., 2020b).

Once the cell commits to spreading, it generally expands its contact footprint in an isotropic, essentially concentric fashion, resulting in a more or less round contact region throughout the spreading phase. This behavior is fundamentally different from migratory cell motions like chemotaxis, which require sustained cell polarization not only in terms of morphology and mechanics, but also intracellular signaling. It seems logical to attribute this difference to the nature of the stimulus, which is uniform in one case and localized in the other. However, especially our substrates coated with low densities of IgG are unlikely to present perfectly uniform IgG layers. Yet even on these substrates, the cell roundness was unaffected by the IgG density, providing further support to the “all-or-nothing” notion of phagocytic spreading. This result also makes sense biologically, considering that a phagocyte should be able to engulf pathogens that are not uniformly coated with antibodies. Finally, our conclusion that phagocytic spreading tends to proceed in a symmetric fashion is particularly impactful for theoretical investigations, as it reinforces the validity of predictions made by mathematical models of phagocytosis that assume an axisymmetric cell-target configuration.

That said, we did observe larger centroid displacements and a significantly higher directional persistence of cells spreading on lower densities of IgG, in agreement with observations of neutrophils spreading on different densities of fibronectin or BSA (Henry et al., 2014) as well as other cell types, for instance, endothelial cells that spread more unevenly on lower densities of Arg-Gly-Asp (RGD) (Reinhart-King et al., 2005). This behavior appears to resemble haptokinesis, albeit on a very small scale considering that the overall cell displacements rarely exceeded 1 μm.

Our picture of phagocytic spreading would be incomplete without considering its limits. What determines the maximum extent of spreading? Key to answering this question is the mechanical resistance to expansion of the apparent cell surface area, i.e., the cortical tension. This ever-present tension maintains the spherical shape of quiescent immune cells, and it rises as the cell surface area increases during shape changes (Herant et al., 2005; Lee et al., 2011; Lee et al., 2015; Roberts et al., 2020a). Geometrical arguments similar to those leading to Eq. (1) show that the apparent cell surface area grows monotonically as the cell-substrate contact region increases. Thus, the protrusive force generated by the cytoskeleton meets with increasing mechanical resistance during cell spreading. The magnitude of the outward pushing force is not unlimited, and neither is the amount of material that can be recruited to the cell surface to enable its further growth. Either one of these closely related constraints will stall spreading. Remarkably, our estimate of the maximum apparent cell surface area produced during frustrated phagocytic spreading agrees well with values measured previously in various types of experiments (Hallett and Dewitt, 2007; Herant et al., 2005; Lee et al., 2015; Simon and Schmid-Schonbein, 1988).

Even before the cell-substrate contact region reaches its maximum extent, the rising resistance due to the cortical tension slows spreading. A higher substrate density of adhesive IgG ligands is expected to aid further spreading at this stage, because the distance over which the protruding cell membrane needs to be displaced before reaching the next adhesion site is smaller. The observed dependence of the maximum extent of spreading on the IgG density (Fig. 5E) confirms that this is indeed the case. Yet, this dependence is much less pronounced than predicted by the Brownian zipper model. This finding underlines the role of adhesion as an important facilitator of spreading, regulating the cell’s commitment to spreading, the maximum contact area, and the directional movement of the contact region. However, the results of this and other studies leave little doubt that active cellular protrusion rather than adhesion is the primary driving force of phagocytic spreading.

## Materials and methods

### Surface preparation

After partial assembly of the 3D-printed chamber (Fig. 2A), the surface of the glass coverslip serving as chamber bottom was incubated with 1% (10 mg/ml) bovine serum albumin (BSA; VWR, Radnor, PA) in phosphate buffered saline (PBS; Bio-Rad, Hercules, CA) for 1 hour at room temperature, then rinsed with PBS. The surface was then incubated with a mixture of polyclonal rabbit anti-BSA IgG (Cat# A11133; Invitrogen, Waltham, MA) and monoclonal mouse anti-BSA IgG-1 (Cat# MA182941, Invitrogen, Waltham, MA) for another hour at room temperature. All mixtures contained a total of 150 µg/ml IgG, which was sufficient to saturate the surface, but the ratio of rabbit to mouse IgG concentration was varied to produce solutions containing 0%, 0.1%, 1%, 10%, or 100% rabbit IgG. After a final rinse with Hanks’ balanced salt solution (HBSS with Ca^2+^ and Mg^2+^; Thermo Fisher Scientific, Waltham, MA), the chamber was loaded with HBSS containing 2% human serum albumin (HSA; MP Biomedicals, Irvine, CA) to block any uncoated regions.

To quantify the actual density of rabbit IgG on coated glass surfaces, coverslips of each batch were incubated with 30 µg/ml anti-human IgG Fc (cross-reacts with rabbit) conjugated to Alexa Fluor-488 (Cat# 409321; BioLegend, San Diego, CA) for 45 minutes at room temperature in the dark, then rinsed with PBS before imaging. Beads from the Quantum Simply Cellular Kit (QSC; Bangs Laboratories, Fishers, IN) were incubated with the same secondary antibody at 6.25 µg/ml, again for 45 minutes at room temperature in the dark, then washed 3 times in PBS with 0.01% Tween and deposited onto plain glass coverslips for imaging.

### Quantification of IgG surface density from fluorescence microscopy

As outlined in the main text (Fig. 1), we determined the density of deposited rabbit IgG by comparing the fluorescence intensities of labeled coverslips to a calibration curve created using a set of QSC beads with known numbers of antibody-binding sites. All fluorescence images were taken on a spinning disc confocal microscope (488 nm laser, 100x oil objective, NA = 1.4). Due to the large spread of fluorescence intensities of the inspected surfaces, we used two different exposure times *t*_*e*_ = 1 s and *t*_*e*_ = 2 s and adjusted the originally measured intensities *I* to a corrected value *Ī* using

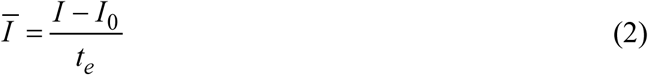

where *I*_*0*_ is the background intensity, i.e., the intensity at the substrate surface in the absence of fluorophore. To minimize photobleaching, we first preadjusted the microscope focus at a suitable location, then moved the microscope stage to a neighboring field of view and acquired a *Z*-stack of images of this unbleached region.

In fluorescence measurements of coverslips, we chose a region of interest of at least 10,000 pixels and determined the most frequent pixel intensity in this region for each image using kernel density estimation. After plotting the most frequent intensity values of all images of a given stack as a function of *Z*-height and fitting a cubic polynomial to these data, we calculated the maximum of the polynomial fit and identified its function value as the fluorescence intensity of the plane containing the fluorophore.

A more elaborate procedure was needed to measure the pertinent fluorescence intensity of labeled QSC beads. We reasoned that the fluorescence at the underside of a bead resting at the chamber bottom is most closely relatable to the fluorescence at the top surface of labeled coverslips. Accordingly, we measured for each selected bead the fluorescence intensity at the center of all bead images in a given Z-stack and then again used polynomial fits to determine the interpolated maximum value of these intensities, which we identified as the fluorescence intensity corresponding to the known density of fluorophore coating the bead. Details of this procedure are illustrated in Fig. S1. Having quantified the relationship between the surface density of rabbit IgG and the fluorescence intensity in this manner, we were able to establish the number of rabbit IgG molecules per μm^2^ of the surface of functionalized coverslips for each of the used IgG incubation mixtures, i.e., for each nominal concentration of rabbit IgG (Fig. 1).

### Cell isolation

The protocol for this study was approved by the Institutional Review Board of the University of California, Davis and all donors provided written informed consent. Human neutrophils were isolated from whole blood of healthy donors by immunomagnetic negative selection using the EasySep Direct Human Neutrophil Isolation Kit (STEMCELL Technologies, Vancouver, Canada). Isolation began from an initial blood volume of 0.5-1 ml, and was carried out according to the protocol described by the manufacturer. Immediately following isolation, the cells were resuspended at 5-10×10^6^ cells per ml in HBSS without Ca^2+^ and Mg^2+^ (Thermo Fisher Scientific) to ensure their quiescence prior to stimulation by contact with IgG.

### Contact area analysis

The cell-substrate contact area was quantified using customized MATLAB software based on code graciously provided by Daniel Kovari (Kovari et al., 2016). In brief, the RICM image was thresholded on both intensity and variance to construct an initial binary image by detecting regions where the local variance was greater than σ^2^_min_ or the intensity was less than *I*_max_ (both thresholds could be adjusted on an image-by-image basis). The binary image was then convolved with a Gaussian kernel to smooth the outline of the cell footprint. Holes in the footprint were filled to define the total contact area. In a few cases where the automatic recognition algorithm performed poorly (especially during early spreading), the cell outlines were traced manually.

The spreading speed was quantified using sigmoidal fits to contact-area-versus-time graphs whenever possible. When necessary, a linear fit was used over a smaller range of the contact area data. The maximum contact area was determined as the average over the plateau region of the curve. To calculate the roundness of the cell-substrate contact region, we determined the largest inscribed and the smallest circumscribed circle of the cell footprint using open-source MATLAB functions (D’Errico, 2017; Warrier, 2017) (Fig. 6A) and took the ratio the radii of these two circles.

### Analysis of centroid directional persistence

We restricted our analysis of the directionality of the motion of the geometric centroid of the substrate contact region of each cell to the phase of fastest contact-area growth, where the spreading speed was greater than 0.4 µm^2^/s. To assess for how long the centroid motion typically maintained a preferred direction, we evaluated the directional autocorrelation function (Gorelik and Gautreau, 2014). First, for each pair of successive video frames (recorded at times *t*_*i*_ and *t*_*i*+1_, respectively), we determined the angle θ(*t*_*i*_) between the *x*-axis and the line of centroid displacement using

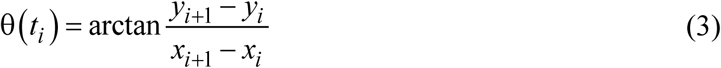

Then, after identifying all pairs of angles θ(*t*_*i*_) and θ(*t*_*j*_) along a given centroid trajectory for which the time gap *t*_*j*_ – *t*_*i*_ equaled a given value Δ*t*, we calculated the autocorrelation function AC(Δ*t*) defined by

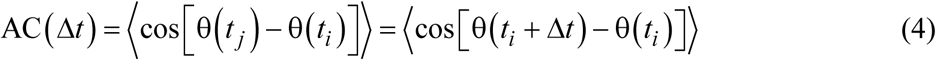

where the angular brackets ⟨…⟩ are used to denote the average over all such pairs. Persistent times are typically defined as the time constants of exponentially decaying autocorrelation functions; however, because of a limited number of data points in this analysis, the autocorrelation functions of the centroids of individual cells often did not lend themselves to exponential fits. Instead, recalling that the persistence time of an exponential decay equals the area under the autocorrelation curve, we here defined the directional persistence time by

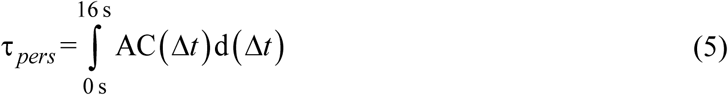

### Computational modeling

Details of our computer model, including all equations and parameter values, are presented in our companion paper (Francis and Heinrich, 2022). In short, we represent the cell interior as a highly viscous Newtonian fluid and assume mechanical equilibrium. This allows us to solve a perturbed form of the Stokes equations at each time step using the finite element method, yielding the flow profile inside the cell. In order to complete this calculation, the relevant stresses are computed for a given shape of the cell, comprising *(i)* adhesion stress, assuming an attractive potential acting between the cell membrane and the IgG-coated surface, *(ii)* protrusion stress, assuming this stress acts normal to the cell membrane and decays exponentially along the membrane, and *(iii)* cortical stress, assuming a uniform cortical tension whose magnitude depends on the amount of surface area expansion as previously quantified for human neutrophils (Herant et al., 2005). All calculations are carried out in MATLAB; our code is available at https://github.com/emmetfrancis/phagocyticSpreading.

### Statistics

All tests for significance were conducted using one-way ANOVA followed by Tukey’s post-hoc test in Origin software. Values more than 1.5 interquartile ranges above the upper quartile or below the lower quartile were excluded as outliers prior to ANOVA.

## Acknowledgements

We thank Dr Soichiro Yamada for his help with confocal microscopy, Dr Daniel Kovari for providing MATLAB code for the analysis of the cell-substrate contact regions, and Dr Robert Guy for many helpful conversations regarding our computational model of cell spreading.

## Competing interests

The authors declare no competing or financial interests.

## Funding

This work was supported by grant R01 GM098060 from the National Institutes of Health, USA, National Science Foundation Graduate Research Fellowship, award number 1650042, and a graduate student research award from the ARCS Foundation.

## Notes

### Competing Interest Statement

The authors have declared no competing interest.

### Summary of Updates

Added Supplemental material.

## References

Allen, L. A. and Aderem, A. (1996). Molecular definition of distinct cytoskeletal structures involved in complement- and Fc receptor-mediated phagocytosis in macrophages. J Exp Med 184, 627–37.

Barger, S. R., Reilly, N. S., Shutova, M. S., Li, Q., Maiuri, P., Heddleston, J. M., Mooseker, M. S., Flavell, R. A., Svitkina, T., Oakes, P. W. et al. (2019). Membrane-cytoskeletal crosstalk mediated by myosin-I regulates adhesion turnover during phagocytosis. Nat Commun 10, 1249.

Ben M’Barek, K., Molino, D., Quignard, S., Plamont, M. A., Chen, Y., Chavrier, P. and Fattaccioli, J. (2015). Phagocytosis of immunoglobulin-coated emulsion droplets. Biomaterials 51, 270–277.

Bournazos, S., Wang, T. T. and Ravetch, J. V. (2016). The Role and Function of Fcgamma Receptors on Myeloid Cells. Microbiol Spectr 4.

Cannon, G. J. and Swanson, J. A. (1992). The macrophage capacity for phagocytosis. J Cell Sci 101 (Pt 4), 907–13.

Chamaraux, F., Fache, S., Bruckert, F. and Fourcade, B. (2005). Kinetics of cell spreading. Phys Rev Lett 94, 158102.

Champion, J. A. and Mitragotri, S. (2006). Role of target geometry in phagocytosis. Proc Natl Acad Sci U S A 103, 4930–4.

Cougoule, C., Wiedemann, A., Lim, J. and Caron, E. (2004). Phagocytosis, an alternative model system for the study of cell adhesion. Semin Cell Dev Biol 15, 679–89.

Cuvelier, D., Thery, M., Chu, Y. S., Dufour, S., Thiery, J. P., Bornens, M., Nassoy, P. and Mahadevan, L. (2007). The universal dynamics of cell spreading. Curr Biol 17, 694–9.

D’Errico, J. (2017). A suite of minimal bounding objects. MATLAB Central File Exchange.

Dewitt, S. and Hallett, M. B. (2002). Cytosolic free Ca(2+) changes and calpain activation are required for beta integrin-accelerated phagocytosis by human neutrophils. J Cell Biol 159, 181–189.

Dewitt, S. and Hallett, M. B. (2020). Calpain Activation by Ca(2+) and Its Role in Phagocytosis. Adv Exp Med Biol 1246, 129–151.

DiNapoli, K. T., Robinson, D. N. and Iglesias, P. A. (2021). A mesoscale mechanical model of cellular interactions. Biophys J 120, 4905–4917.

Dobereiner, H. G., Dubin-Thaler, B., Giannone, G., Xenias, H. S. and Sheetz, M. P. (2004). Dynamic phase transitions in cell spreading. Phys Rev Lett 93, 108105.

Dubin-Thaler, B. J., Giannone, G., Dobereiner, H. G. and Sheetz, M. P. (2004). Nanometer analysis of cell spreading on matrix-coated surfaces reveals two distinct cell states and STEPs. Biophys J 86, 1794–806.

Dubin-Thaler, B. J., Hofman, J. M., Cai, Y., Xenias, H., Spielman, I., Shneidman, A. V., David, L. A., Dobereiner, H. G., Wiggins, C. H. and Sheetz, M. P. (2008). Quantification of cell edge velocities and traction forces reveals distinct motility modules during cell spreading. PLoS One 3, e3735.

Dustin, M. L. (2016). Complement Receptors in Myeloid Cell Adhesion and Phagocytosis. Microbiol Spectr 4.

Etienne, J. and Duperray, A. (2011). Initial dynamics of cell spreading are governed by dissipation in the actin cortex. Biophys J 101, 611–21.

Fardin, M. A., Rossier, O. M., Rangamani, P., Avigan, P. D., Gauthier, N. C., Vonnegut, W., Mathur, A., Hone, J., Iyengar, R. and Sheetz, M. P. (2010). Cell spreading as a hydrodynamic process. Soft Matter 6, 4788–4799.

Flannagan, R. S., Jaumouille, V. and Grinstein, S. (2012). The cell biology of phagocytosis. Annu Rev Pathol 7, 61–98.

Footer, M. J., Kerssemakers, J. W. J., Theriot, J. A. and Dogterom, M. (2007). Direct measurement of force generation by actin filament polymerization using an optical trap. Proceedings of the National Academy of Sciences of the United States of America 104, 2181–2186.

Francis, E. A. and Heinrich, V. (2022). Integrative experimental/computational approach establishes active cellular protrusion as the primary driving force of phagocytic spreading by immune cells. (submitted).

Freeman, S. A. and Grinstein, S. (2014). Phagocytosis: receptors, signal integration, and the cytoskeleton. Immunol Rev 262, 193–215.

Frisch, T. and Thoumine, O. (2002). Predicting the kinetics of cell spreading. J Biomech 35, 1137–41.

Futosi, K., Fodor, S. and Mocsai, A. (2013). Neutrophil cell surface receptors and their intracellular signal transduction pathways. Int Immunopharmacol 17, 638–50.

Gorelik, R. and Gautreau, A. (2014). Quantitative and unbiased analysis of directional persistence in cell migration. Nat Protoc 9, 1931–43.

Hallett, M. B. and Dewitt, S. (2007). Ironing out the wrinkles of neutrophil phagocytosis. Trends Cell Biol 17, 209–14.

Hategan, A., Sengupta, K., Kahn, S., Sackmann, E. and Discher, D. E. (2004). Topographical pattern dynamics in passive adhesion of cell membranes. Biophys J 87, 3547–60.

Heinrich, V. (2015). Controlled One-on-One Encounters between Immune Cells and Microbes Reveal Mechanisms of Phagocytosis. Biophys J 109, 469–76.

Henry, S. J., Crocker, J. C. and Hammer, D. A. (2014). Ligand density elicits a phenotypic switch in human neutrophils. Integr Biol (Camb) 6, 348–56.

Herant, M., Heinrich, V. and Dembo, M. (2005). Mechanics of neutrophil phagocytosis: behavior of the cortical tension. J Cell Sci 118, 1789–97.

Herant, M., Heinrich, V. and Dembo, M. (2006). Mechanics of neutrophil phagocytosis: experiments and quantitative models. J Cell Sci 119, 1903–13.

Herant, M., Lee, C. Y., Dembo, M. and Heinrich, V. (2011). Protrusive push versus enveloping embrace: computational model of phagocytosis predicts key regulatory role of cytoskeletal membrane anchors. PLoS Comput Biol 7, e1001068.

Janmey, P. A., Hinz, B. and McCulloch, C. A. (2021). Physics and Physiology of Cell Spreading in Two and Three Dimensions. Physiology (Bethesda) 36, 382–391.

Jaumouille, V., Cartagena-Rivera, A. X. and Waterman, C. M. (2019). Coupling of beta2 integrins to actin by a mechanosensitive molecular clutch drives complement receptor-mediated phagocytosis. Nat Cell Biol 21, 1357–1369.

Jaumouille, V. and Waterman, C. M. (2020). Physical Constraints and Forces Involved in Phagocytosis. Front Immunol 11, 1097.

Kovar, D. R. and Pollard, T. D. (2004). Insertional assembly of actin filament barbed ends in association with formins produces piconewton forces. Proc Natl Acad Sci U S A 101, 14725–30.

Kovari, D. T., Wei, W., Chang, P., Toro, J. S., Beach, R. F., Chambers, D., Porter, K., Koo, D. and Curtis, J. E. (2016). Frustrated Phagocytic Spreading of J774A-1 Macrophages Ends in Myosin II-Dependent Contraction. Biophys J 111, 2698–2710.

Lam, J., Herant, M., Dembo, M. and Heinrich, V. (2009). Baseline mechanical characterization of J774 macrophages. Biophys J 96, 248–54.

Lee, C. Y., Herant, M. and Heinrich, V. (2011). Target-specific mechanics of phagocytosis: protrusive neutrophil response to zymosan differs from the uptake of antibody-tagged pathogens. J Cell Sci 124, 1106–14.

Lee, C. Y., Thompson, G. R., 3rd, Hastey, C. J., Hodge, G. C., Lunetta, J. M., Pappagianis, D. and Heinrich, V. (2015). Coccidioides Endospores and Spherules Draw Strong Chemotactic, Adhesive, and Phagocytic Responses by Individual Human Neutrophils. PLoS One 10, e0129522.

Lim, J. J., Grinstein, S. and Roth, Z. (2017). Diversity and Versatility of Phagocytosis: Roles in Innate Immunity, Tissue Remodeling, and Homeostasis. Front Cell Infect Microbiol 7, 191.

Lubeck, M. D., Steplewski, Z., Baglia, F., Klein, M. H., Dorrington, K. J. and Koprowski, H. (1985). The interaction of murine IgG subclass proteins with human monocyte Fc receptors. J Immunol 135, 1299–304.

Nelsen, E., Hobson, C. M., Kern, M. E., Hsiao, J. P., O’Brien Iii, E. T., Watanabe, T., Condon, B. M., Boyce, M., Grinstein, S., Hahn, K. M. et al. (2020). Combined Atomic Force Microscope and Volumetric Light Sheet System for Correlative Force and Fluorescence Mechanobiology Studies. Sci Rep 10, 8133.

Odenthal, T., Smeets, B., Van Liedekerke, P., Tijskens, E., Van Oosterwyck, H. and Ramon, H. (2013). Analysis of initial cell spreading using mechanistic contact formulations for a deformable cell model. PLoS Comput Biol 9, e1003267.

Ostrowski, P. P., Freeman, S. A., Fairn, G. and Grinstein, S. (2019). Dynamic Podosome-Like Structures in Nascent Phagosomes Are Coordinated by Phosphoinositides. Dev Cell 50, 397–410 e3.

Pacheco, P., White, D. and Sulchek, T. (2013). Effects of microparticle size and Fc density on macrophage phagocytosis. PLoS One 8, e60989.

Parekh, S. H., Chaudhuri, O., Theriot, J. A. and Fletcher, D. A. (2005). Loading history determines the velocity of actin-network growth. Nat Cell Biol 7, 1219–23.

Paul, D., Achouri, S., Yoon, Y. Z., Herre, J., Bryant, C. E. and Cicuta, P. (2013). Phagocytosis dynamics depends on target shape. Biophys J 105, 1143–50.

Reinhart-King, C. A., Dembo, M. and Hammer, D. A. (2005). The dynamics and mechanics of endothelial cell spreading. Biophys J 89, 676–89.

Richards, D. M. and Endres, R. G. (2014). The mechanism of phagocytosis: two stages of engulfment. Biophys J 107, 1542–53.

Roberts, R. E., Dewitt, S. and Hallett, M. B. (2020a). Membrane Tension and the Role of Ezrin During Phagocytosis. Adv Exp Med Biol 1246, 83–102.

Roberts, R. E. and Hallett, M. B. (2019). Neutrophil Cell Shape Change: Mechanism and Signalling during Cell Spreading and Phagocytosis. Int J Mol Sci 20.

Roberts, R. E., Martin, M., Marion, S., Elumalai, G. L., Lewis, K. and Hallett, M. B. (2020b). Ca(2+)-activated cleavage of ezrin visualised dynamically in living myeloid cells during cell surface area expansion. J Cell Sci 133.

Rosales, C. and Uribe-Querol, E. (2017). Phagocytosis: A Fundamental Process in Immunity. Biomed Res Int 2017, 9042851.

Shashidharamurthy, R. B. E.; Patel, J.; Kaur, R.; Meganathan, J.; Selvaraj, P. (2010). Analysis of cross-species IgG binding to human and mouse Fcgamma receptors (FcγRs). The Journal of Immunology 184.

Simon, S. I. and Schmid-Schonbein, G. W. (1988). Biophysical aspects of microsphere engulfment by human neutrophils. Biophys J 53, 163–73.

Tran-Son-Tay, R., Needham, D., Yeung, A. and Hochmuth, R. M. (1991). Time-dependent recovery of passive neutrophils after large deformation. Biophys J 60, 856–66.

Uribe-Querol, E. and Rosales, C. (2020). Phagocytosis: Our Current Understanding of a Universal Biological Process. Front Immunol 11, 1066.

van Zon, J. S., Tzircotis, G., Caron, E. and Howard, M. (2009). A mechanical bottleneck explains the variation in cup growth during FcgammaR phagocytosis. Mol Syst Biol 5, 298.

Vorselen, D., Barger, S. R., Wang, Y., Cai, W., Theriot, J. A., Gauthier, N. C. and Krendel, M. (2021). Phagocytic ‘teeth’ and myosin-II ‘jaw’ power target constriction during phagocytosis. Elife 10.

Vorselen, D., Wang, Y., de Jesus, M. M., Shah, P. K., Footer, M. J., Huse, M., Cai, W. and Theriot, J. A. (2020). Microparticle traction force microscopy reveals subcellular force exertion patterns in immune cell-target interactions. Nat Commun 11, 20.

Warmerdam, P. A., van de Winkel, J. G., Gosselin, E. J. and Capel, P. J. (1990). Molecular basis for a polymorphism of human Fc gamma receptor II (CD32). J Exp Med 172, 19–25.

Warrier, R. (2017). max_inscribed_circle. MATLAB Central File Exchange.

Wolfenson, H., Iskratsch, T. and Sheetz, M. P. (2014). Early events in cell spreading as a model for quantitative analysis of biomechanical events. Biophys J 107, 2508–14.

Xiong, Y., Rangamani, P., Fardin, M. A., Lipshtat, A., Dubin-Thaler, B., Rossier, O., Sheetz, M. P. and Iyengar, R. (2010). Mechanisms controlling cell size and shape during isotropic cell spreading. Biophys J 98, 2136–46.

Yeung, A. and Evans, E. (1989). Cortical shell-liquid core model for passive flow of liquid-like spherical cells into micropipets. Biophys J 56, 139–49.

Zhang, Y., Hoppe, A. D. and Swanson, J. A. (2010). Coordination of Fc receptor signaling regulates cellular commitment to phagocytosis. Proc Natl Acad Sci U S A 107, 19332–7.

