## Supplemental Figure, Figure Legend, and Movie Legend for "Mechanisms of frustrated phagocytic spreading of human neutrophils on antibody-coated surfaces"

**Supplementary information**

**Figure legends**

**Fig. S1. Procedure to quantify the fluorescence at the underside of labeled beads.** (**A**) Confocal image of QSC beads labeled with secondary antibody resting at the bottom of a chamber. A search box (*yellow square*) was placed around each isolated bead. (**B**) Robust determination of the center of bead images. For each image, we first computed axes of mirror symmetry in a user-definable number of directions. We used 12 such axes in the present analysis (*orange lines*). For each direction, the symmetry axis was determined by shifting a straight line across the region of interest containing the bead image. For each position of this line, we calculated the cross-correlation of the bead image with its respective mirror image obtained by flipping the original image about this line. The line position corresponding to the interpolated maximum of cross-correlation values then defined the symmetry axis in the given direction. Finally, the bead center was calculated as the center of mass of the cross-sections of all pairs of symmetry axes. This procedure gave consistent results even for moderately noisy bead images and bead images with dark center regions. (**C**) Filmstrip representation of a *Z*-stack of confocal images of a QSC bead. The focal plane was moved from below the bead's underside (image (1)) to a height where it cut through the bead slightly above its lowest point (image (13)). The center of each bead image in the stack was determined as explained in (B). (**D**) Circularly averaged radial intensity profile of a confocal bead image. We calculated the mean intensities along circles centered about the center of the bead image, spaced out by one pixel (*blue symbols*). We then fit an even, low-order polynomial function to a subrange of the discrete intensity data (*red curve*). We identified the fluorescence intensity at the center of the bead image with the function value of the fitting function at radius zero. (**E**) Plot of the polynomial fitting functions obtained for bead images of the stack of (C). For clarity, only the graphs for every other bead image are shown. (**F**) Determination of the fluorescence intensity at the lowest point of the bead surface. The intensity values at the center of all bead images of the stack of (C) (obtained as explained in (D)) are plotted as a function of their number within the stack (*blue symbols*). We fit a low-order polynomial function to a subrange of these data (*orange curve*) and identified the sought fluorescence intensity at the underside of the bead with the calculated maximum of the polynomial fitting function

**Movie legends**

**Movie S1. Examples of human neutrophils spreading on antibody-coated surfaces.** Three timelapse image sequences show neutrophils (initially apparent as blurry bright spots) approaching and settling onto the bottom coverslip. The cells then proceed to spread on the IgG-coated surface. The quickly growing cell-substrate contact area of each cell appears as a dark footprint in the RICM images. The first sequence illustrates the general behavior of the neutrophils, but our analysis only included isolated cells that were spreading without obstruction by other cells, such as shown in the second sequence. The third sequence focuses on a single, enlarged cell footprint and displays the cell over an extended period of time to also illustrate its more or less random motion during the post-spreading phase.

**Figure S1**

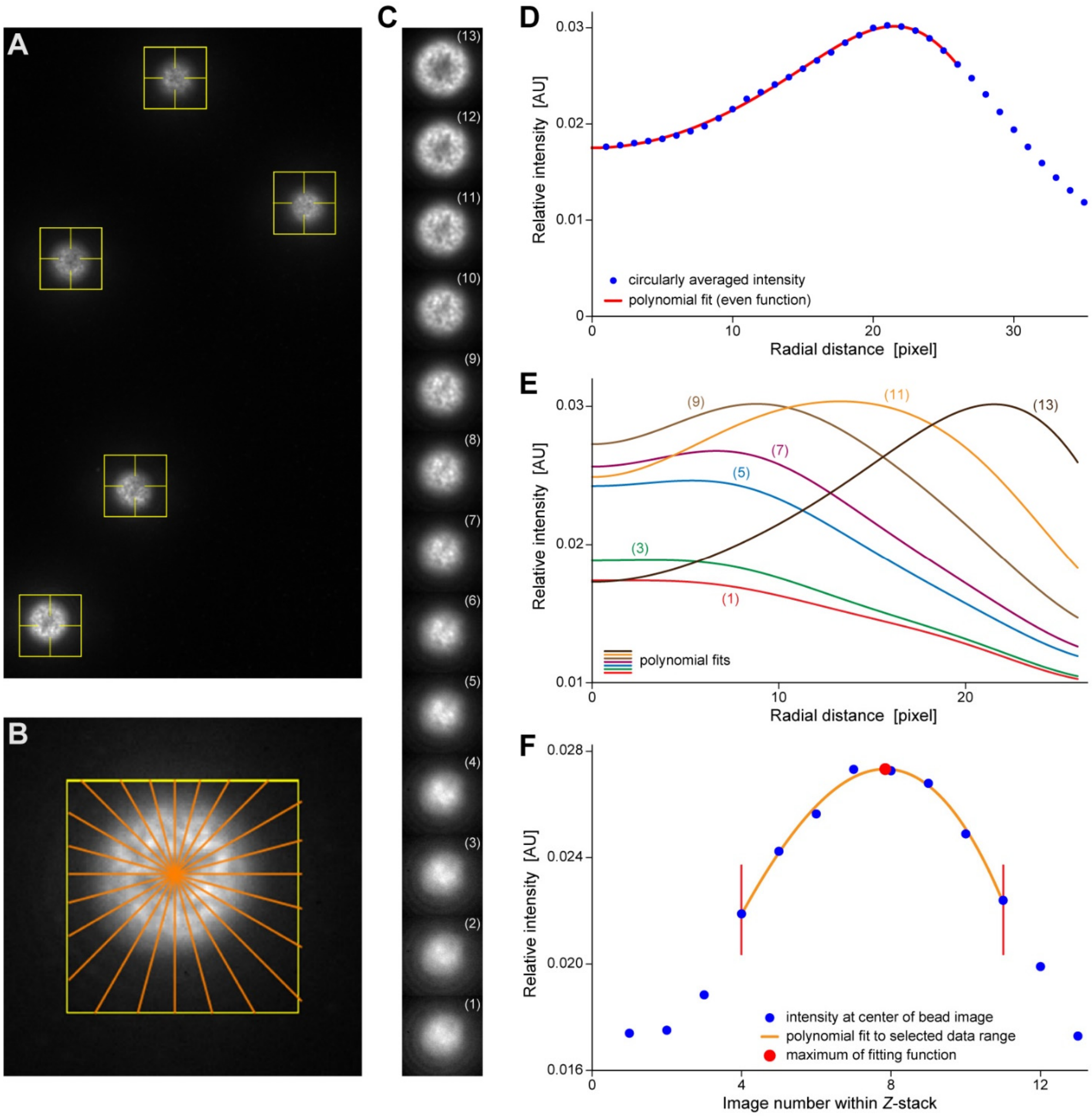

A
B
C
(13)
(12)
(11)
(10)
(9)
(8)
(7)
(6)
(5)
(4)
(3)
(2)
(1)
D
Relative intensity [AU]
circularly averaged intensity
polynomial fit (even function)
Radial distance [pixel]
E
Relative intensity [AU]
polynomial fits
Radial distance [pixel]
F
Relative intensity [AU]
intensity at center of bead image
polynomial fit to selected data range
maximum of fitting function
Image number within Z-stack
